# In culture cross-linking of bacterial cells reveals proteome scale dynamic protein-protein interactions at the peptide level

**DOI:** 10.1101/094961

**Authors:** Luitzen de Jong, Edward A. de Koning, Winfried Roseboom, Hansuk Buncherd, Martin J. Wanner, Irena Dapic, Petra J. Jansen, Jan H. van Maarseveen, Garry L. Corthals, Peter J. Lewis, Leendert W. Hamoen, Chris G. de Koster

**Affiliations:** Swammerdam Institute for Life Sciences, 1098 XH Amsterdam, The Netherlands; Van’t Hoff Institute of Molecular Science, University of Amsterdam, 1098 XH Amsterdam, The Netherlands.; Faculty of Medical Technology, Prince of Songkla University, Hatyai, Songkhla 90110, Thailand.; School of Environmental and Life Sciences, University of Newcastle, Callaghan, NSW 2308, Australia.

## Abstract

Identification of dynamic protein-protein interactions at the peptide level on a proteomic scale is a challenging approach that is still in its infancy. We have developed a system to cross-link cells directly in culture with the special lysine cross-linker bis(succinimidyl)-3-azidomethyl-glutarate (BAMG). We used the Gram positive model bacterium *Bacillus subtilis* as an exemplar system. Within 5 min extensive intracellular cross-linking was detected, while intracellular cross-linking in a Gram-negative species, *Escherichia coli*, was still undetectable after 30 min, in agreement with the low permeability in this organism for lipophilic compounds like BAMG. We were able to identify 82 unique inter-protein cross-linked peptides with less than a 1% false discovery rate by mass spectrometry and genome-wide data base searching. Nearly 60% of the inter-protein cross-links occur in assemblies involved in transcription and translation. Several of these interactions are new, and we identified a binding site between the δ and β′ subunit of RNA polymerase close to the downstream DNA channel, providing a clue into how δ might regulate promoter selectivity and promote RNA polymerase recycling. Our methodology opens new avenues to investigate the functional dynamic organization of complex protein assemblies involved in bacterial growth.

## INTRODUCTION

Understanding how biological assemblies function at the molecular level requires knowledge of the spatial arrangement of their composite proteins. Chemical protein cross-linking coupled to identification of proteolytic cross-linked peptides by mass spectrometry (CX-MS) has been successfully used to obtain information about the 3D topology of isolated protein complexes (1). In this approach, the amino acid sequences of a cross-linked peptide pair reveal the interacting protein domains. The continued increase in peptide identification sensitivity by improved MS techniques and equipment has opened the door to proteome-wide protein interaction studies by cross-linking living cells. Such a systems-level view on dynamic protein interactions would be a tremendously powerful tool to study cell biology.

The only large scale *in vivo* CX-MS studies with bacteria thus far have been performed with four Gram-negative species and have provided valuable data about the 3D topology of outer membrane and periplasmic protein complexes (2-5). However, with less than 1% of the total dataset of cross-linked peptides, the fraction of detected interactions between known soluble cytoplasmic proteins was underrepresented (2,3), possibly caused by poor membrane permeability of the relatively large cross-linker used in these studies, by the barrier formed by the outer membrane and periplasmic space, or by repeated washing and buffer exchange steps before cross-linking that may have led to dissociation of transient interactions. We also noted biological inconsistencies in these studies as several tens of cross-links revealed interactions between soluble cytoplasmic proteins and outer membrane proteins (3) suggesting some cell lysis occurred during the harvesting/washing stages prior to cross-linking. These results show that identification of *bone fide* interactions between cytoplasmic proteins is a non-trivial issue that has not yet been satisfactorily solved.

Here we describe a system that fulfills all requirements for efficient trapping and identification of both stable and dynamic protein-protein interactions in the cytoplasm of living cells. We used the Gram positive model *Bacillus subtilis*, widely studied for processes guided by dynamic protein-protein interactions involved in gene expression, cell division, sporulation and germination (6). Cross-linking was accomplished by a previously designed reagent, bis(succinimidyl)-3-azidomethyl-glutarate (BAMG) (Figure 1, Supporting Information)(7). This bifunctional N-hydroxysuccinimidyl ester covalently links juxtaposed lysine residues on protein surfaces *via* two amide bonds bridged by a spacer of 5 carbon atoms. The relatively short spacer results in high-resolution cross-link maps. A cross-linker with the same spacer length and similar hydrophobicity, disuccinimidyl glutarate (DSG) (Figure 1, Supporting Information), is membrane permeable and has been used before for cross-linking in living human cells (8). Importantly, to prevent dissociation of transient intracellular protein interactions by washing and medium change, we added the cross-linker directly in the culture medium containing a low concentration of primary amines to minimize reaction with and quenching of the cross-linker.

**Figure 1.**
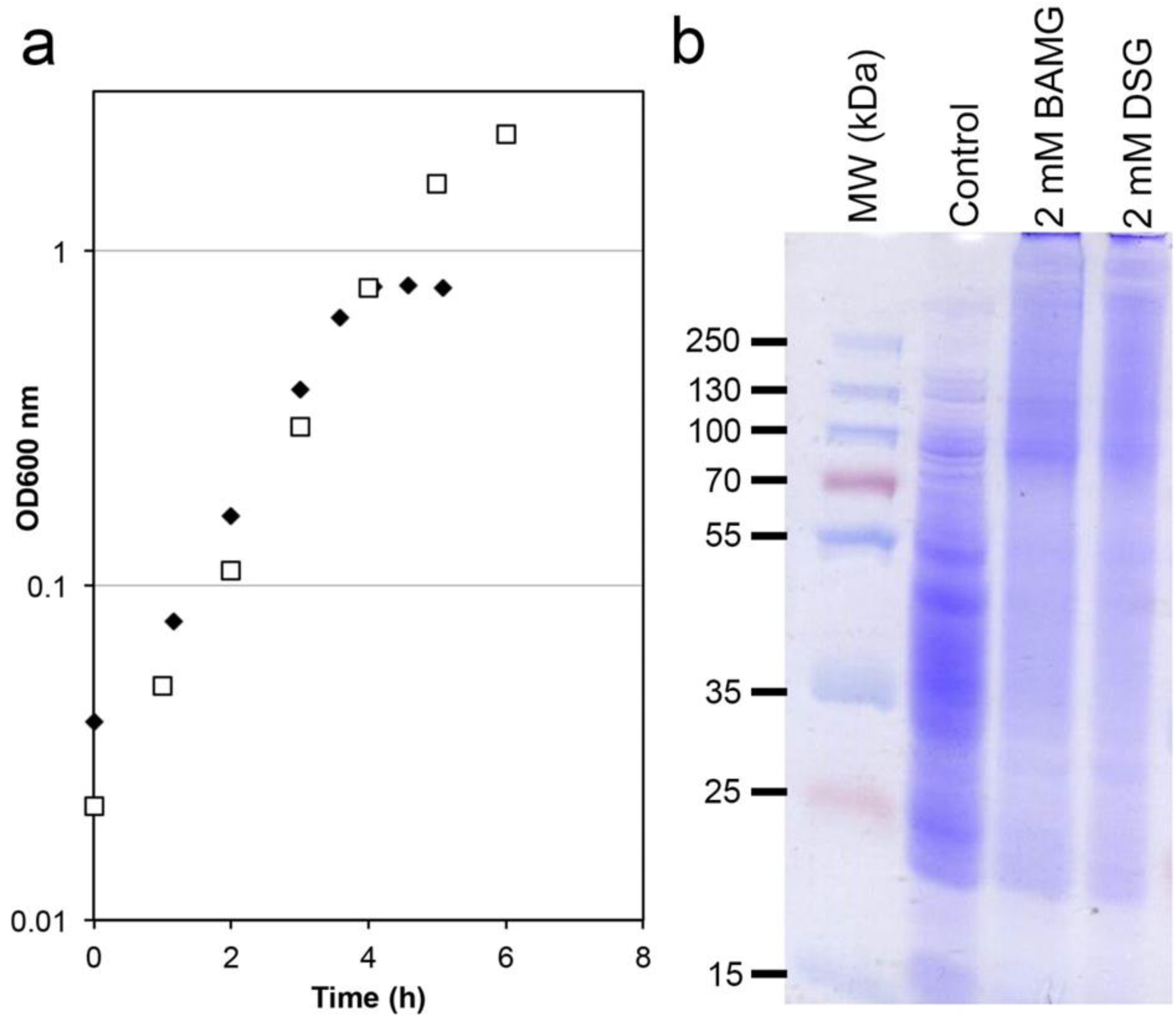
In vivo cross-linking in of *B. subtilis* in culture. **a**, Growth curves of *B. subtilis* in minimal medium with 1.2 mM glutamine (filled diamonds) and 5 mM glutamine (open squares); **b**, SDS-PAGE analysis of in vivo cross-linking with BAMG and DSG of exponentially growing B. subtilis directly in the growth medium. Control, soluble proteins from untreated exponentially growing B. subtilis. 2 mM BAMG or DSG, soluble proteins from exponentially growing B. subtilis treated with 2 mM BAMG or DSG, respectively. Molecular weights (kDa) are shown on the left hand side adjacent to pre-stained molecular weight markers (MW markers).

A main limitation of proteome-wide cross-linking studies is the identification of cross-linked peptides in total cell extracts. This is facilitated both by separation of cross-linked peptides from the bulk of unmodified species and by determination of the masses of the two linked peptides. To this end BAMG provides the cross-linked peptides with additional chemical properties that greatly facilitate cross-link identification by virtue of the presence of a 3-azidomethylene group in the spacer domain. The azido group can be reduced to an amine group, enabling isolation of the low abundant cross-linked peptides by two-dimensional strong cation exchange chromatography (9). In addition, chemical reduction renders the two cross-link amide bonds of BAMG-cross-linked peptides scissile in the gas phase by collision-induced dissociation, in a way that the masses of the two composite peptides can be determined from an MS/MS spectrum, thereby facilitating peptide identification by searching an entire genomic database (10). Identification of cross-linked peptides from a single MS/MS spectrum provides BAMG with a large advantage over the two other cleavable reagents used up to now that require multi-stage tandem mass spectrometry to map cross-links formed *in vivo* (3,11). To obtain sufficient cross-linked material by labeling directly in culture, adequate amounts of cross-linker are necessary, and in this report we also include a scalable new synthesis route for BAMG.

Using our novel *in vivo* crosslinking procedure, we were able to detect several transient protein-protein interactions at the peptide level in *B. subtilis* cells. Many of the inter-protein crosslinks could be corroborated by structural data from previous studies, but other crosslinks represent new interactions. A cross-link revealing the binding site between the β’ and δ subunits of RNA polymerase (RNAP) demonstrates the power of *in vivo* cross-linking directly in the culture medium to obtain insight into the molecular mechanisms of action of complex dynamic assemblies active in growing cells. This approach can be readily modified to allow the identification of less abundant protein complexes or to investigate in depth the dynamic assembly of specific protein complexes.

## EXPERIMENTAL PROCEDURES

### Synthesis of BAMG

**Scheme 1.**
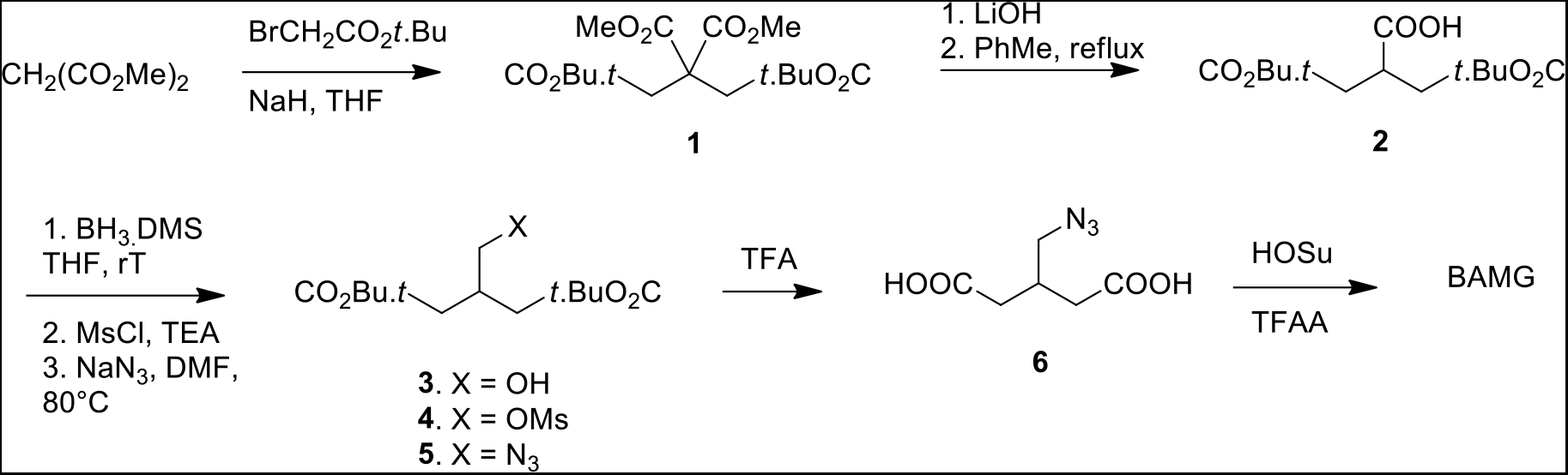
Formation of BAMG.

#### Tetra-ester **1**

Dimethyl malonate (3.42 ml, 30 mmol) was added dropwise to a stirred suspension of sodium hydride (60% dispersion in oil, 2.46 g, 66.0 mmol) in THF (120 ml) at RT. After stirring for 45 minutes *tert*. butyl bromoacetate (9.45 ml, 64 mmol) was added dropwise. The reaction was stirred for 16 h, cooled in ice, and the excess sodium hydride was carefully neutralized with acetic acid (ca 6 mmol). Extractive workup with sat. aquous NH_4_Cl and ether, drying over MgSO_4_ and evaporation gave tetra-ester **1** as an oil (quantitative) which was immediately used for the next step. ^1^H-NMR (400 MHz, CDCl_3_): 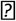3.77 (s, 6H); 3.06 (s, 4H); 1.45 (s, 18H).

#### Carboxylic acid **2**

A solution of tetra-ester **1** (30 mmol) in THF (150 ml) and methanol (40 ml) was diluted with a solution of lithium hydroxide (2.94 g, 70 mmol) in water (150 ml) and refluxed for 2 h. After removal of the organic solvents *in vacuo* the aqueous layer was extracted with a 1 : 1 mixture of diethyl ether and PE 40/60. Acidification of the water layer (pH ca 1), extraction with diethyl ether, drying with MgSO_4_ and evaporation gave a mixture of mono-and di-carboxylic acids. This mixture was refluxed in toluene (150 ml) for 2 h. Evaporation of the toluene gave carboxylic acid **2** (5.5 gr, 19.1 mmol, 64% from dimethyl malonate) as a slowly solidifying oil. ^1^H-NMR (400 MHz, CDCl_3_): 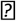3.23 (m, 1H); 2.68 (dd, 1H, *J* = 7.2, *J* = 16.6 Hz); 2.54 (dd, 1H, *J* = 6.2, *J* = 16.6 Hz); 1.46 (s, 18H). ^13^C-NMR (100 MHz, CDCl_3_): 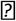179.7, 170.4, 81.1, 37.5, 36.2, 27.8; IR (film, cm^−1^): 3200, 1728, 1711 cm^−1^.

#### Alcohol **3**

Borane dimethylsulfide (1.45 ml, 15 mmol) was added dropwise to a solution of carboxylic acid **2** (1.3 g, 4.5 mmol) in anhydrous THF (30 ml) at 0 °C. The reaction was stirred at RT for 16 h and carefully quenched with saturated aqueous NH_4_Cl and diethyl ether. Extractive workup and flash chromatography with a mixture of PE 40/60 and ethyl acetate (3 : 1 an 1 : 1) gave alcohol **3** (0.78 gr, 2.85 mmol, 63%) as an oil. ^1^H-NMR (400 MHz, CDCl_3_): 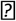3.65 (d, 2H, *J* = 5.2 Hz); 2.45 (m, 1H); 2.3 – 2.4 (m, 4H); 1.47 (s, 18JH); ^13^C-NMR (100 MHz, CDCl_3_): 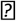172.1, 80.6, 80.55, 65.0, 64.9, 37.2, 37.1, 34.8, 27.9; IR: 3500, 1726 cm^−1^. IR (film, cm^−1^): 3500, 1726.

#### Mesylate **4**

Methanesulfonyl chloride (0.255 ml, 3.3 mmol) was added dropwise to a solution of alcohol **3** (0.78 g, 2.85 mmol) and triethylamine (0.526 ml, 4.0 ml) in anhydrous dichloromethane (20 ml) at 0 °C. After stirring for 1 h at 0 °C the reaction was diluted with diethyl ether (ca 50 ml) and quenched with water. Extractive workup gave mesylate **4** (1.0 g, 2.84 mmol, quantitative). ^1^H-NMR (400 MHz, CDCl_3_): 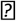4.31 (d, 2H, *J* = 5.1 Hz); 2.67 (m, 1H); 2.35 – 2.47 (m, 4H); 1.47 (m, 18H); ^13^C-NMR (100 MHz, CDCl_3_): 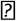170.6, 80.9, 80.85, 71.1, 36.95, 35.9, 31.9, 27.9; IR (film, cm^−1^): 1725.

#### Azide **5**

A mixture of mesylate **4** (1.0 g, 2.84 mmol) and sodium azide (0.554 g, 8.5 mmol) in anhydrous DMF (10 ml) was stirred at 85 °C for 3 h. Extractive workup with water and diethyl ether, followed by chromatography (PE 40/60 : ethyl acetate 6 : 1) gave pure azide **5** (0.84 g, 2.8 mmol, 98% from **3**). ^1^H-NMR (400 MHz, CDCl_3_): 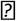3.44 (d, 2H, *J* = 5.7 Hz); 2.50 (m, 1H); 2.3 – .45 (m, 4H); 1.48 (s, 18H); ^13^C-NMR (100 MHz, CDCl_3_): 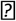170.8, 80.6, 54.1, 32.4, 27.9; IR (film, cm^−1^): 2102, 1728.

#### 3-(Azidomethyl)-glutaric acid **6**

Azide **5** (0.638 g, 2.13 mmol) was stirred in a mixture of dichloromethane (16 ml) and trifluoroacetic acid (4 ml) for 6 h at RT. Toluene (30 ml) was added and the solvents were removed *in vacuo*. Drying of the resulting glass (0.02 mbar, 50 °C) gave pure di-acid **6** in quantitative yield. ^1^H-NMR (400 MHz, CDCl_3_+ 10% CD_3_OD): 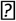3.44 (d, 2H, *J* = 5.7 Hz); ^13^C-NMR (100 MHz, CDCl_3_+ 10% CD_3_OD): 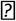176.2, 54.0, 35.7, 31.8; IR (film, cm^−1^): 3100 (broad), 2103, 1708.

#### **BAMG**: Bis(succinimidyl) 3-azidomethyl-glutarate

This step was carried out according to a described procedure (12) Trifluoroacetic anhydride (1.4 ml) was added to a solution of di-acid **6** (0.415 g, 2.1 mmol) and *N*-hydroxysuccinimide (1.15 g, 10.0 mmol) in a mixture of dichloromethane (8 ml) and anhydrous pyridine (4 ml) at 0 °C. The cooling bath was removed and stirring was continued for 1.5 h. The reaction mixture was diluted with dichloromethane, and extracted with three 50 ml portions of 1M HCl and finally with NaHCO_3_(2 × 50 ml). Drying over MgSO_4_, evaporation of the solvent and drying (0.02 mbar, 40 °C) gave BAMG (0.76 g, 2.0 mmol, 95%) as a slightly yellow syrup. BAMG was stored at -80 °C. Before storage, BAMG was dissolved in acetonitrile, divided in aliquots and dried by vacuum centrifugation. ^1^H-NMR (400 MHz, CDCl_3_): 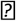3.65 (d, 2H, *J* = 5.5 Hz); 2.83 – 2.90 (m, 12H), 2.75 (m, 1H); ^13^C-NMR (100 MHz,CDCl_3_): 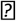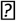169.0, 166.6, 52.5, 32.3, 32.2, 25.4; IR (film, cm^−1^): 2108, 18,14, 1783, 1735.

### Growth of bacteria

*B. subtilis* strain 168 (trp^−^) was grown in a MOPS minimal medium (13) modified as described for *B. subtilis* (14) and supplemented with 0.2% glucose, 1.2 mM glutamine and 0.2 mM tryptophan. To obtain an exponentially growing culture for cross-linking, streaks from a glycerol stock of cells grown on liquid LB medium were first put on an LB agar plate. Following overnight growth at 37°C a single colony was suspended in 10 ml minimal medium in 100 ml culture flasks. From the suspension, dilutions were made into 10 ml minimal medium for overnight growth in 100 ml flasks placed at 37°C in a water bath shaking at 240 rpm. An overnight culture in mid exponential growth as determined by an OD_600 nm_= 0.3-0.5 was used for dilution to OD_600 nm_= 0.01 in pre-warmed minimal medium in Erlenmeyer flasks to obtain exponentially growing cultures for cross-linking. *Escherichia coli* strain MC4100 was cultured in MOPS medium supplemented with 0.16% (w/v) glucosamine and 0.1 mM NH_4_Cl. To obtain an exponentially growing culture for cross-linking, an overnight culture in this growth medium was 40 times diluted in fresh medium to an OD_600 nm_= 0.08.

### Cross-linking ***in vivo***

In exponentially growing *B. subtilis* cultures at OD _600 nm_= 0.45 – 0.50, cross-linking was started by the addition of 2.0 mM BAMG from a freshly prepared stock solution of 1 M in DMSO. A magnetic stirrer was used for rapid mixing with the culture. Cross-linking was for 5 min in the shaking water bath at 37°C. The cross-linking reaction was quenched by the addition of 1M Tris-Cl (pH 8.0) to a final concentration of 50 mM. Cross-linked cells were harvested by centrifugation for 5 min at 4000 g and cell pellets were stored frozen at -20°C.

Exponentially growing *Escherichia coli* cells were cross-linked at OD _600 nm_= 0.7, corresponding to 0.21 μg dry weight per ml (15), with 2 mM BAMG for 5, 10 and 30 min. The cross-linking reaction was quenched by the addition of 50 mM Tris-Cl (pH 8.0). We assess that 2.76 mM glucose, formed from an equivalent amount of N-acetylglucosamine (16), has been consumed for energy and biomass production at the time of cross-linking (13). Deamination of glucosamine, after deacetylation of N-acetylglucosamine, results in the formation of an equivalent amount of NH_4_^+^ of which 1.88 mM NH_4_^+^ has been consumed for biomass production (13). This implies that the culture contains 0.88 mM free NH_4_^+^ that can react with BAMG, leaving 1.12 mM BAMG for protein cross-linking. Cross-linked cells were harvested by centrifugation for 5 min at 4000 g and cell pellets were stored frozen at -20°C.

### Protein extraction

Frozen *B. subtilis* cell pellets from 2-40 ml culture medium were resuspended in 1 ml of a solution containing 10 mM Tris-HCl and 0.1 mM EDTA (pH 7.5). Cell suspensions of 1 ml in 2 ml propylene Eppendorf vials placed in ice water were lysed by sonication with a micro tip, mounted in an MSE ultrasonic integrator operated at 21 Hz and amplitude setting 3, in 6 periods of 15 s with 15 s intervals in between. Lysates were centrifuged for 15 min at 16000 g. Supernatants were used for further analysis. Cell extracts from *Escherichia coli* were prepared by suspending cell pellets from 1 ml cultures in SDS-PAGE sample buffer (17) without β-mercaptoethanol. Suspensions were incubated for 1 h at 60°C and then centrifuged for 2 min at 13000 g. Proteins were concentrated with 0.5 ml Amicon Ultra 10 kDa cut off centrifugal filters (Millipore) before SDS-PAGE analysis.

### Gel filtration

A cross-linked protein fraction with a size distribution of approx. 400 kDa to 1-2 MDa was obtained by gel filtration on a Superose 6 10/300 GL column (GE Healthcare) operated on an Akta FPLC system (GE Healthcare) in a buffer containing 20 mM HEPES pH 7.9, 300 mM KCl, 0.2 mM EDTA, 0.1 mM DTT and 20% glycerol (gel filtration buffer) at a flow rate of 0.5 ml min^−1^. Fractions of 1 ml were collected and snap frozen in liquid nitrogen for storage at −20°C.

### Protein determination and polyacrylamide gel electrophoresis in the presence of sodium dodecyl sulphate (SDS-PAGE)

Protein was measured with the bicinchoninic acid method (18) using a protein assay kit (Pierce). SDS-PAGE (17) was carried out using 10% or 12% precast Novex gels (Themo Fisher Scientific).

### Protein digestion

Pooled gel filtration fractions of extracted cross-linked proteins in the 400 kDa to 1-2 MDa range were concentrated to about 10 mg protein/ml with 0.5 ml Amicon Ultra 10 kDa cut off centrifugal filters (Millipore). Prior to digestion, cysteines were alkylated by addition of a solution of 0.8 M iodoacetamide (Sigma–Aldrich), followed by the addition of solution of 1 M Tris-HCl pH 8.0, 9.6 M urea (Bioreagent grade, Sigma–Aldrich) to obtain final concentrations of 40 mM iodoacetamide, 0.1 M Tris HCl and 6 M urea, respectively. Incubation was for 30 min at room temperature in the dark. The solution was diluted 6 times by the addition of 0.1 M Tris–HCl pH 8.0 and digested with trypsin (Trypsin Gold, Promega, Madison, WI, USA) overnight at 30°C at a 1:50 (w/w) ratio of enzyme and substrate. Peptides were desalted on C18 reversed phase TT3 top tips (Glygen, Columbia, USA), eluted with 0.1% TFA in 50% acetonitrile and dried in a vacuum centrifuge.

### Diagonal SCX chromatography

A protocol described earlier (9) was used with several modifications. The main difference was the use of a solution of ammonium formate instead of KCl for salt gradient elution. The use of the volatile ammonium formate avoids time-consuming desalting steps and prevents loss of material. Dry desalted peptides (240 μg) were reconstituted with 10 μl of a solution containing 0.1% TFA and 25% acetonitrile followed by the addition of 0.2 ml 10 mM ammonium formate and 25% acetonitrile pH 3.0 (buffer A) and 0.2 ml of the mixture was loaded on a Poly-sulfoethyl aspartamide column (2.1 mm ID, 10 cm length) (Poly LC Inc., Columbia, USA) operated on an Ultimate HPLC system (LC Packings, Amsterdam, The Netherlands). For elution, at a flow rate of 0.4 ml min^−1^, increasing amounts of buffer B (500 mM ammonium formate pH 3.0) were mixed with buffer A, according to the following scheme. At t = 5 min, 1% buffer B was added, at t = 10 min a linear gradient from 1% to 50% buffer B was started over 10 min, followed by a gradient from 50% to 100% buffer B over 3 min. Elution with 100% B lasted 2 min after which the column was washed with buffer A for 19 min. A UV detector was used to measure absorbance at 280 nm of the eluent. Peptides started to elute at t = 14 min and were manually collected in 0.2 ml fractions and lyophilized. For secondary SCX runs, dried cross-linked enriched peptides (fractions 7-16^6^) were dissolved in 20 μl 40 mM TCEP (BioVectra) in 20% acetonitrile and incubated under argon for 2 h at 60°C. The peptide solution was then diluted with 0.19 ml buffer A just before loading for the secondary SCX runs. Elution occurred under the same conditions as in the primary SCX run. Material was collected when the absorbance at 280 nm started to rise again (about 30s after the end of the elution time window of the primary fraction, until it came back to base level (high salt shifted fraction). Collected eluent was lyophilized.

### LC-MS/MS

Identification of proteins by LC-MS/MS analysis of peptides in SCX fractions with an AmaZon Speed Iontrap with a CaptiveSpray ion source (Bruker) coupled with an EASY-nLC II chromatographic system (Proxeon,Thermo Scientific) and data processing have been described in detail before (19).

Identification of cross-linked peptides enriched by diagonal SCX chromatography by LC-MS/MS analysis was performed with an Eksigent Expert nanoLC 425 system connected to the Nano spray source of a TripleTOF 5600+ mass spectrometer. Peptides were loaded onto an Eksigent trap column (Nano LC trap set, ChromXP C18, 120 Å, 350 μm × 0.5 mm) in a solution containing 0.1 % TFA, and 2 % acetonitrile and desalted with 3% TFA and 0.1 % formic acid at 2 μL/min. After loading, peptides were separated on an in-house packed 7 cm long, 75 μm inner diameter analytical column (Magic C18 resin, 100 Å pore size, 5 μm) at 300 nL/min. Mobile phase A consisted of 0.1% formic acid in water and mobile phase B consisted of 0.1 % formic acid in acetonitrile. The gradient consisted of 5% B for 5 min, then 5-10 % B over 10 min, followed by 10-35 % B over 60 min and then the gradient was constant at 80 % B for 10 min. After each run the column was equilibrated for 20 min at starting conditions. The TripleTOF 5600+ mass spectrometer was operated with nebulizer gas of 6 PSI, curtain gas of 30 PSI, an ion spray voltage of 2.4 kV and an interface temperature of 150°C. The instrument was operated in high sensitivity mode. For information-dependent acquisition, survey scans were acquired in 50 ms in the m/z range 400-1250 Da. In each cycle, 20 product ion scans were collected for 50 ms in the m/z range 100-1800 Da, if exceeding 100 counts per seconds and if the charge state was 3+ to 5+. Dynamic exclusion was used for half of the peak width (15s) and rolling collision energy was used.

Before acquisition of two samples the mass spectrometer was calibrated using the built-in autocalibration function of Analyst 1.7. For MS calibration, 25 fmol of β-galactosidase digest (Sciex) was injected. For TOF MS calibration, ions with the following m/z values were selected: 433.88, 450.70, 528.93, 550.28, 607.86, 671.34, 714.85 and 729.40 Da. The ion at m/z 729.4 Da was selected for fragmentation and product ions were used for TOF MS/MS calibration.

For 27 out of 29 LC-MS/MS runs, average mass deviations from calculated values of identified components varied from -4.0 ± 2.4 to 15.3 ± 4.1 ppm. For data processing of MS/MS (MS1MS2) data by Reang (described below) and database searching of MS/MS (MS1MS2) data by Mascot, 25 ppm mass tolerance was allowed in these cases for both M1 and MS2. In the two remaining runs average mass deviations of identified components were 31.9 ± 12.0 and 62.5 ± 7.8 ppm, respectively. In these cases, a mass tolerances of 50 ppm and 75 ppm, respectively, was allowed for both MS1 and MS2

### Data processing

Raw LC-MS1MS2 data were processed with Mascot Distiller and MS2 data were deconvoluted to MH^+^ values at the QStar default settings using the option to calculate masses for 3+ to 6+ charged precursor ions in case the charge state could not be assessed unambiguously.

### Identification of candidate cross-linked peptides

For cross-link identification using the entire *B. subtilis* sequence database, a software tool named Reang (10) was used for further MS1MS2 data processing. The rationale of the processing by Reang described below is based on the notion that an MS1MS2 spectrum of BAMG-cross-linked peptides provides both the information for the masses of the candidate composing peptides as well as the fragment ions for identification of the composing peptides. In brief, Reang identifies precursor ions with mass > 1500 Da, potentially corresponding to a BAMG-cross-linked peptide pair A and B with the azide reduced to an amine, showing evidence for cleavage of the cross-linked amide bonds in the presumed cross-link. Such cleavage events result in product ions of the unmodified peptides A and B and in modified peptides Am and Bm fulfilling the following mass relationships

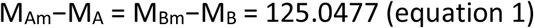
EQN1

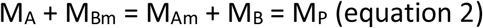
EQN2
where M_Am_ and M_Bm_, resp., are the masses of peptides A and B modified with the remnant m of the cross-linker in the form of a γ-lactam with elemental composition C_6_H_7_NO_2_, corresponding to a mass of 125.0477 Da, M_A_ and M_B_ are the masses of peptide A and peptide B, resp., and M_P_ is the mass of the precursor P.

Reang identifies among the 30 product ions of highest signal intensity within a mass error of 25-75 ppm (i) pairs of mass values of fragment ions > 500 Da differing 125.0477 Da, i.e., a candidate A and Am pair or B and Bm pair, (ii) pairs of mass values for A and B fulfilling the equation M_A_+ M_B_+ 125.0477 = M_P_, and (iii) pairs of mass values for Am and Bm fulfilling the equation M_Am_+ M_Bm_-125.0477 = M_P_. The mass values of the other pairs in the cases (i), (ii) and (iii) are calculated from eq. 1 and eq. 2.

MS1 values of entries in the MS1MS2 data files with MS2 data fulfilling at least one of the equations 1 or 2 are replaced by MS1 values corresponding to M_A_, M_Am_, M_B_ and M_Bm_.

Furthermore, fragment ions corresponding to M_A_, M_Am_, M_B_ and M_Bm_ are removed from the new MS1MS2 entries as well as fragments ions larger than the new MS1 values.

The new MS1 MS2 files in pkl format are input for Mascot to nominate candidate peptides for A, Am, B and Bm by interrogating the *B. subtilis* strain 168 database containing both forward and reversed sequences. Reang combines the nominated peptides with a Mascot score ≥ 1 into candidate cross-linked peptides and assigns these candidates with a mass tolerance of 25-75 ppm to precursor ions in the original MS1 MS2 data file. Candidates are validated based on the original MS1 MS2 data files. The principle of our approach is that an MS1 MS2 spectrum of cross-linked peptides provides both the information for the masses of the candidate composing peptides as well as the fragment ions for identification of the composition of the peptides.

### Cross-link mapping and validation

Validation and false discovery rate determination is facilitated by a software tool, called Yeun Yan (10). Only one candidate cross-linked peptide or cross-linked decoy peptide is assigned for each precursor ion, at least if the candidate fulfills certain criteria with respect to a minimum number of y ions that should be assigned with a mass tolerance of 25-75 ppm to each of the composing peptides in a cross-linked peptide pair. Only assigned y ions among the 100 fragments of highest signal intensity are taken into account. The number of required assigned y ions differs for intra-protein and inter-protein cross-linked peptides, the latter type of cross-links requiring more stringent criteria for assignment than the former type. This difference is based on the notion (10,20-22) that that the probability of identifying cross-links as the result of a random event from a sequence database of many proteins is higher for cross-linked peptides from different protein sequences (inter-protein cross-links) than for cross-linked peptides comprising different peptide sequences from the same protein sequence (intra-protein cross-links). Intra-protein cross-links comprise peptides from the same protein sequence, whereas inter-protein cross-links comprise peptides from different protein sequences, unless the peptides have identical sequences, and, therefore, must have originated from two identical protein molecules in a complex, assuming that a given protein sequence does not yield two or more identical tryptic peptides. In the case of an intra-protein cross-link, at least one unambiguous y ion should be assigned for each composing peptide and both the number of assigned y ions for each composing peptide and the score, called the Yeun Yan score, defined below, should be the same as or more than the number of assigned y ions and the score for other possible candidates with forward sequences or one or more decoy sequences for the same precursor. No intra-protein cross-link decoy sequences consisting of reversed sequences from the same protein or hybrid forward and reversed sequences from the same protein were observed. For an inter-protein cross-linked peptide pair between different proteins or decoy cross-links, the number of assigned y ions should be at least 3 for each peptide built up from up to 10 amino acid residues and at least 4 for peptides consisting of 11 amino acids or more. The number of assigned y ions for each peptide should be the same or more than the number of assigned y ions for each peptide of other possible candidates for the same precursor ion. Both the total number of y ions and the Yeun Yan score for a candidate cross-linked peptide should exceed the total number of y ions and the score for other possible candidates for the same precursor. These criteria are also used for assignment of inter-protein cross-links comprising two identical sequences. For both intra-and inter-protein cross-links, a Yeun Yan score of more than 40 is required. We do not take into account the number of b ions as a requirement for assignment, since b ions in our dataset occur more than four times less than y ions, and taking them into account would require application of different statistical weights for assignment of b and y ions, which would complicate the calculations. Some spectra with a precursor mass difference of +1 Da compared with an identified cross-linked peptide were manually inspected to verify whether the precursor represents a cross-linked peptide in which the azide group was converted by TCEP to a hydroxyl group instead of an amine group (7). This appeared to be the case on a single occasion.

For proposed candidate cross-linked peptides, Yeun Yan calculates the masses of possible b and y fragments, b and y fragments resulting from water loss (b0, y0) and ammonia loss (b*, y*), fragment ions resulting from cleavage of the amide bonds of the cross-link, and b, b0, b*, y, y0 and y* fragments resulting from secondary fragmentations of cleavage products. A prerequisite for nomination by Yeun Yan as a candidate and calculation of the corresponding score is the presence in the MS2 spectrum of at least ten fragment ions and assignment of one unambiguous y ion per peptide. A y ion is considered ambiguous if it can also be assigned to one or more other fragments. A y ion resulting from primary and secondary cleavage at the same position is counted only once for the requirement with respect to the minimal number of unambiguous y ions for validation and assignment.

The YY score is calculated according to the equation

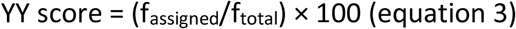
EQN1
in which f_assigned_ is the total number of matching fragment ions, including primary b and y fragments, b and y fragments resulting from water loss (b0, y0) and ammonia loss (b*, y*), fragment ions resulting from cleavage of the amide bonds of the cross-link, and b, b0, b*,y, y0 and y* fragments resulting from secondary fragmentation of products resulting from cross-link amide bond cleavages, and f_total_ is the total number of fragments ions in the spectrum with a maximum of 40, starting from the fragment ion of highest intensity.

### Cross-linking of isolated RNAP

RNAP was purified from a pellet from 2L of culture of *B. subtilis* BS200 (*trpC2 spo0A3 rpoC-6his spc*) as follows. Following lysis in 20 mM KH_2_PO_4_ pH8.0, 500 mM NaCl, 0.1mM DTT and clarification, RNAP was initially purified by Ni^2+^ affinity chromatography. Pooled RNAP containing fractions were dialysed in 20 mM Tris-HCl pH7.8, 150 mM NaCl, 1 mM EDTA, 0.1 mM DTT and loaded onto a MonoQ column(GE Healthcare) in dialysis buffer without EDTA. RNAP was eluted using a gradient over 10 column volumes in dialysis buffer supplemented with 2M NaCl. RNAP containing fractions were pooled and dialysed in 20 mM Tris-HCl pH7.8, 150 mM NaCl, 10 mM MgCl_2_, 30% glycerol, 0.1 mM DTT prior to flash freezing and storage at -80°C. Before cross-linking RNAP was dialyzed in 20 mM HEPES, 150 mM NaCl, 10% glycerol, pH 7.4 (cross-linking buffer). RNAP was cross-linked at a protein concentration of 0.5 mg/ml for 30 min at room temperature. The cross-link reaction was started by the addition of a solution containing 80 mM BAMG in acetonitrile to obtain a final concentration of 0.4 mM BAMG and 0.5% acetonitrile. The reaction was quenched by adding 1 M Tris–HCl pH 8.0 to a final concentration of 50 mM. Digestion of the cross-linked protein and isolation and identification of cross-linked peptides was carried out as described previously (23).

### Determination of spatial distances between cross-linked residues

PDB files of structural models were downloaded from the protein data bank (http://www.rcsb.org/pdb/home/home.do). Only PDB files of *B. subtilis* proteins or proteins with at least 40% sequence identity were used. Sequences were aligned using the BLAST algorithm to identify corresponding residues. (http://blast.ncbi.nlm.nih.gov/Blast.cgi?PAGE=Proteins). For RNAP, a homology model of *B. subtilis* elongation complex (24) (EC) was used. It gives similar results as with PDB file 2O5I (25). Structures were inspected with DeepView - Swiss-PdbViewer ( http://spdbv.vital-it.ch/refs.html ) for distance measurements.

### In silico docking

A homology model of was used along with other published structures identified by their protein data bank IDs (PDB ID) detailed below. The N-terminal domain of δ (PDB ID 2M4K) was used along with the EC model and *in vitro* and *in vivo* cross-linking data to produce a model using the HADDOCK2.2 web server Easy interface (26) 40 models in 10 clusters (4 models per cluster) were obtained and analysed for compliance to the maximum Cα-Cα cross-link distance permitted by BAMG (29.7Å) in PyMol v1.8.2.0. The total cumulative distance of β’K208-δ48, β’K1104-δ48, and β’K1152-δ48 Cα measurements was used to identify models that were most compliant (lowest cumulative distance) with cross-link criteria. To co-localise δ and σ^A^ region 1.1, *E. coli* RNAP holoenzyme in which σ^70^ region 1.1 was present (PDB ID 4LK1) (27) was super-imposed over the *B. subtilis* EC model and all but σ^70^ region 1.1 deleted.

## RESULTS

### Defined growth medium for in vivo cross-linking

Addition of the cross-linker directly to a growth medium enables trapping of transient protein interactions in living cells that may otherwise dissociate upon washing and medium exchange. This requires a low concentration of primary amines to minimize quenching of the cross-linker in the medium. We found that the growth rate of *B. subtilis* in minimal medium containing only 1.2 mM glutamine as the nitrogen source was almost identical to the growth rate using the standard 5 mM glutamine, with doubling times of 45 and 43 min, respectively (Figure 1a). Addition of 2 mM of the cross-linker BAMG resulted in an immediate end to the increase in OD_600 nm_, indicating that biomass production ceased instantaneously (Figure 2, Supporting Information). As shown by SDS-PAGE analysis (Figure 1b), most extracted proteins become cross-linked upon treatment of the cells with 2 mM BAMG for 5 min. The same results were obtained with disuccinimidyl glutarate (DSG) (Figure 1b). This indicates that the azidomethylene group in BAMG does not affect membrane permeability, with DSG and BAMG having about the same protein cross-linking efficiency (7). SDS-PAGE analysis (Figure 3, Supporting Information) shows that the cross-linked proteins could be digested efficiently, establishing a set of experimental conditions suitable for the identification of *in vivo* cross-linked peptides.

**Figure 2.**
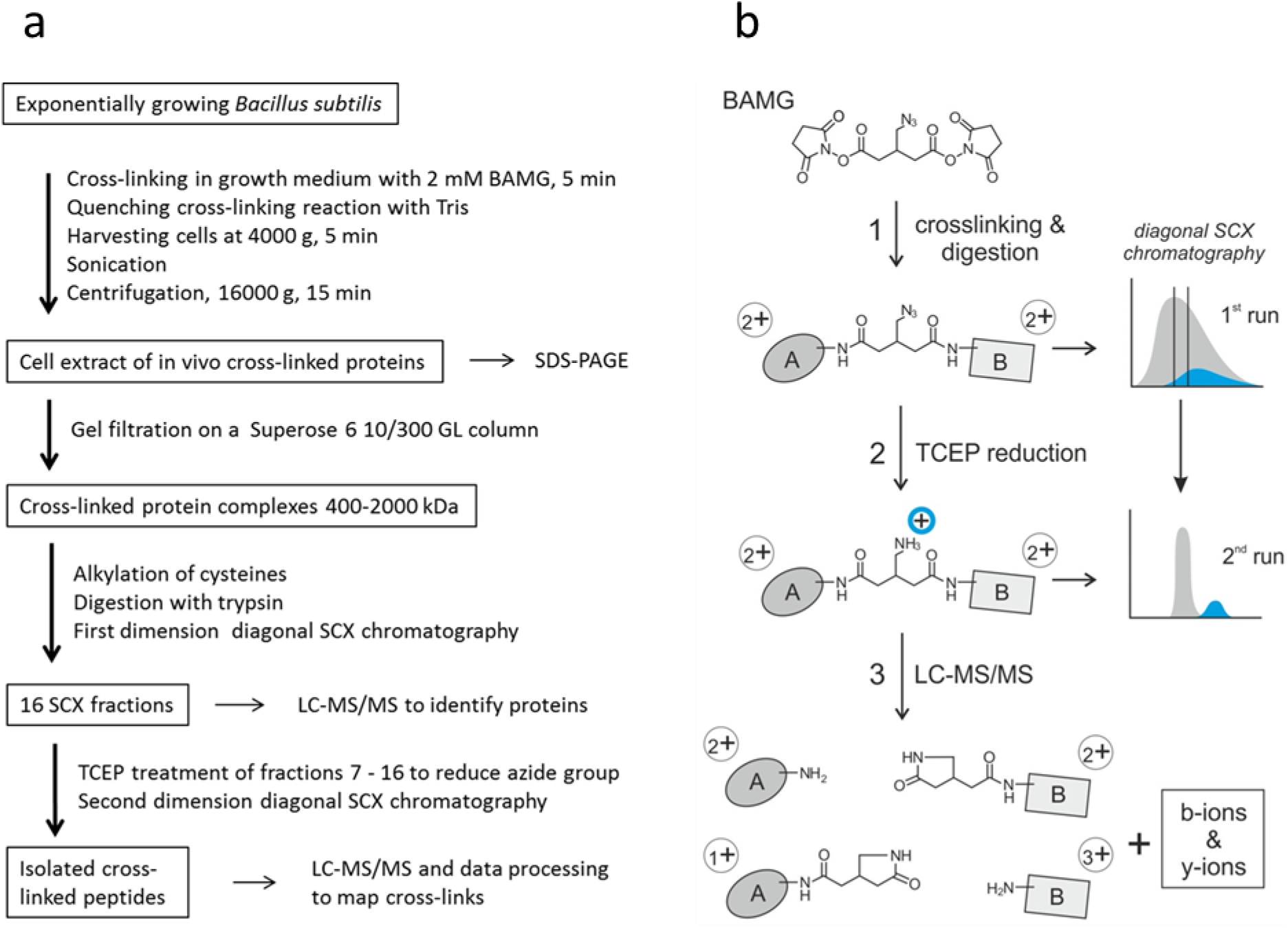
Workflow for peptide level identification of protein cross-links introduced by BAMG in exponentially growing *B. subtilis*. **a**, overview; **b**, left part, reaction products formed (1) in the cross-link reaction with BAMG, (2) by TCEP-induced reduction and (3) by cross-link amide bond cleavages and peptide bond cleavages by collision with gas molecules during LC-MS/MS leading to formation of unmodified peptide ions and peptide ions modified by the cross-linker remnant in the form of a γ-lactam, along with b and y ions. A, peptide A; B, peptide B. Depicted peptide charge states after (1) and (2) are calculated for pH 3, assuming full protonation of basic amino acids and carboxylic acids. Depicted charge states in the gas phase after (3) are arbitrary, assuming a net charge state of +4 of the intact precursor ion. Right part of panel **b**, principles of isolation of cross-linked peptides by diagonal strong cation exchange (DSCX) chromatography. After digestion, the peptide mixture from a protein extract is fractionated by SCX chromatography, using a mobile phase of pH 3 and a salt gradient of ammonium formate to elute bound peptides (1st run). Cyan, cross-linked peptides; grey, unmodified peptides. subsequently, fractions containing cross-linked peptides are treated with TCEP to reduce the azido group to an amine group, which becomes protonated at pH 3, adding one positive charge to cross-linked peptides. TCEP-treated fractions are then separately subjected to a second run of diagonal chromatography. The change in chromatographic behavior caused by the charge increase of cross-linked peptides leads to their separation from the bulk of unmodified peptides present in the same primary SCX fraction.

**Figure 3.**
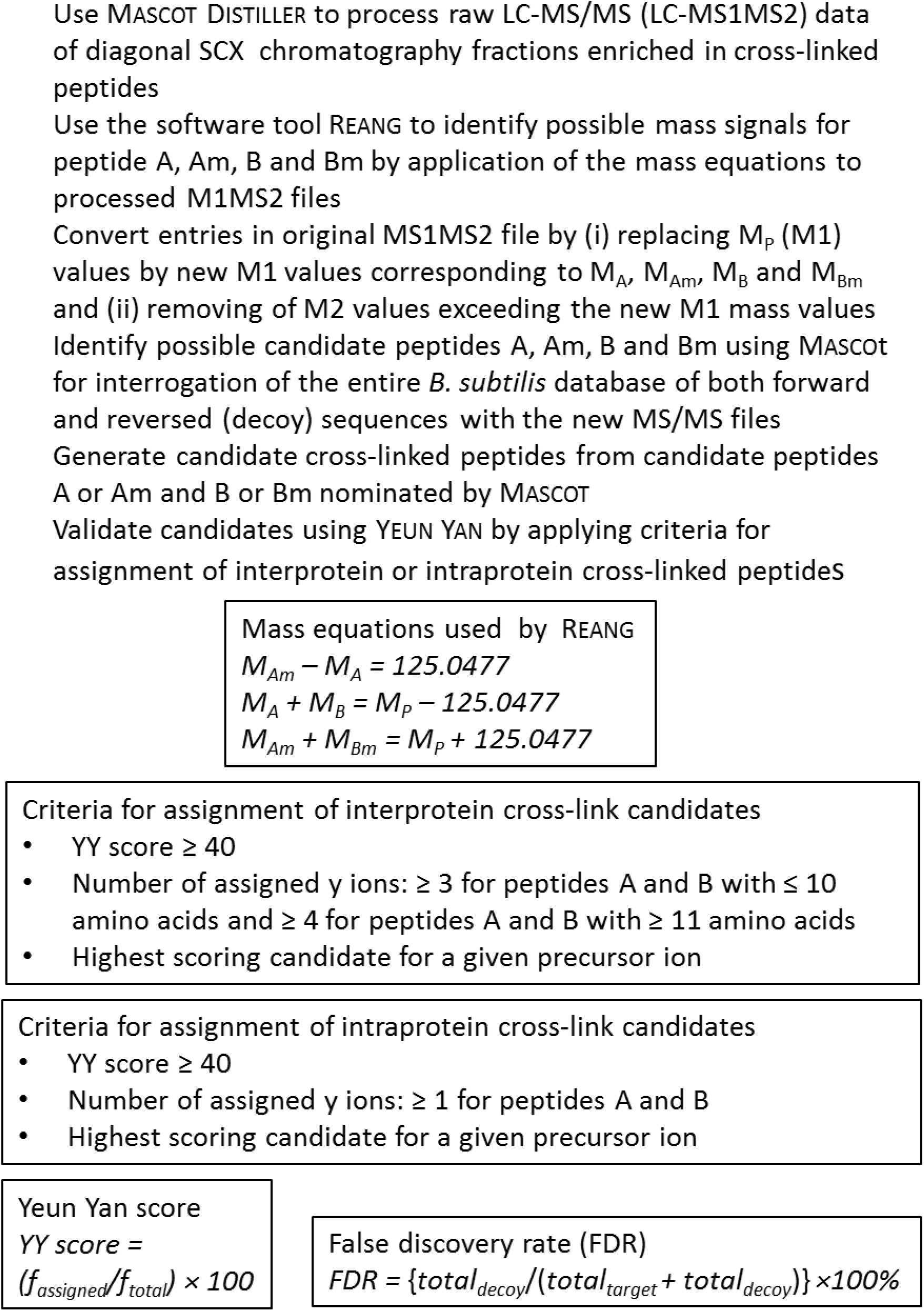
Overview of identification and validation of cross-linked peptides by mass spectrometry and database searching. A, B, Am, Bm, free peptides A and B and peptides A and B modified by the cross-linker in the form of a γ-lactam; M_A_, M_B_, M_Am_, M_Bm_, masses of peptides A and B and their γ-lactam modifications; M_P_, precursor mass; f_assigned_, total number of assigned fragment ions; f_total_, total number of fragment ions of highest intensity taken into account with a minimum of 10 and a maximum of 40 fragments; total_decoy_, total number of assigned decoy peptides; total_target_, total number of assigned target peptides.

### In vivo cross-linking of Gram-negative species

It is well known that the outer membrane of Gram negative bacteria forms a barrier for the diffusion of lipophilic compounds, due to the relatively low fluidity of the bilayer imposed by the lipopolysaccharide outer leaflet (28). To test whether this property prevents the use of BAMG as an effective cross-linker for soluble proteins in Gram negative species, we used *Escherichia coli* as an example. Cells were grown in a MOPS medium with 0.16% N-acetylglucosamine as the only source for energy, carbon and nitrogen. The culture medium was also supplied with 0.1 mM NH_4_Cl to provide the cells with a small amount of a nitrogen source to enable a rapid start of growth. When the NH_4_^+^ in the medium has been consumed, cells have to rely on the ammonia that is formed intracellularly during catabolism of N-acetylglucosamine. Based on published data (13) it can be calculated that the amount of NH_4_^+^ thus formed will be enough for fast growth, and will not accumulate to concentrations that will decrease the concentration of BAMG by more than 1 mM by reaction with the cross-linker (Experimental procedures). The doubling time under these conditions is 56 min during mid exponential growth. This compares favorably with a similar, but slightly faster, doubling time of 50-55 min reported by others using a medium containing N-acetylglucosamine supplemented with 9.5 mM NH_4_Cl as a nitrogen source (16). Upon addition of 2 mM BAMG to the exponentially growing cells in this medium, inhibition of growth occurred (Figure 4, Supporting Information). However, in striking contrast with the *Bacillus subtilis* results (Figure 1b), SDS-PAGE analysis provided no evidence for large scale cross-linking of extracted proteins from BAMG-treated *E. coli* cells, since the Coomassie-blue-stained patterns in the lanes from control cells and cross-linked cells were indistinguishable (Figure 5, Supporting Information). These results are in agreement with the known slow diffusion rate of hydrophobic compounds through the outer membrane and put limitations on the use of Gram-negative organisms for rapid *in vivo* cross-linking by N-hydroxysuccinimidyl esters.

**Figure 4.**
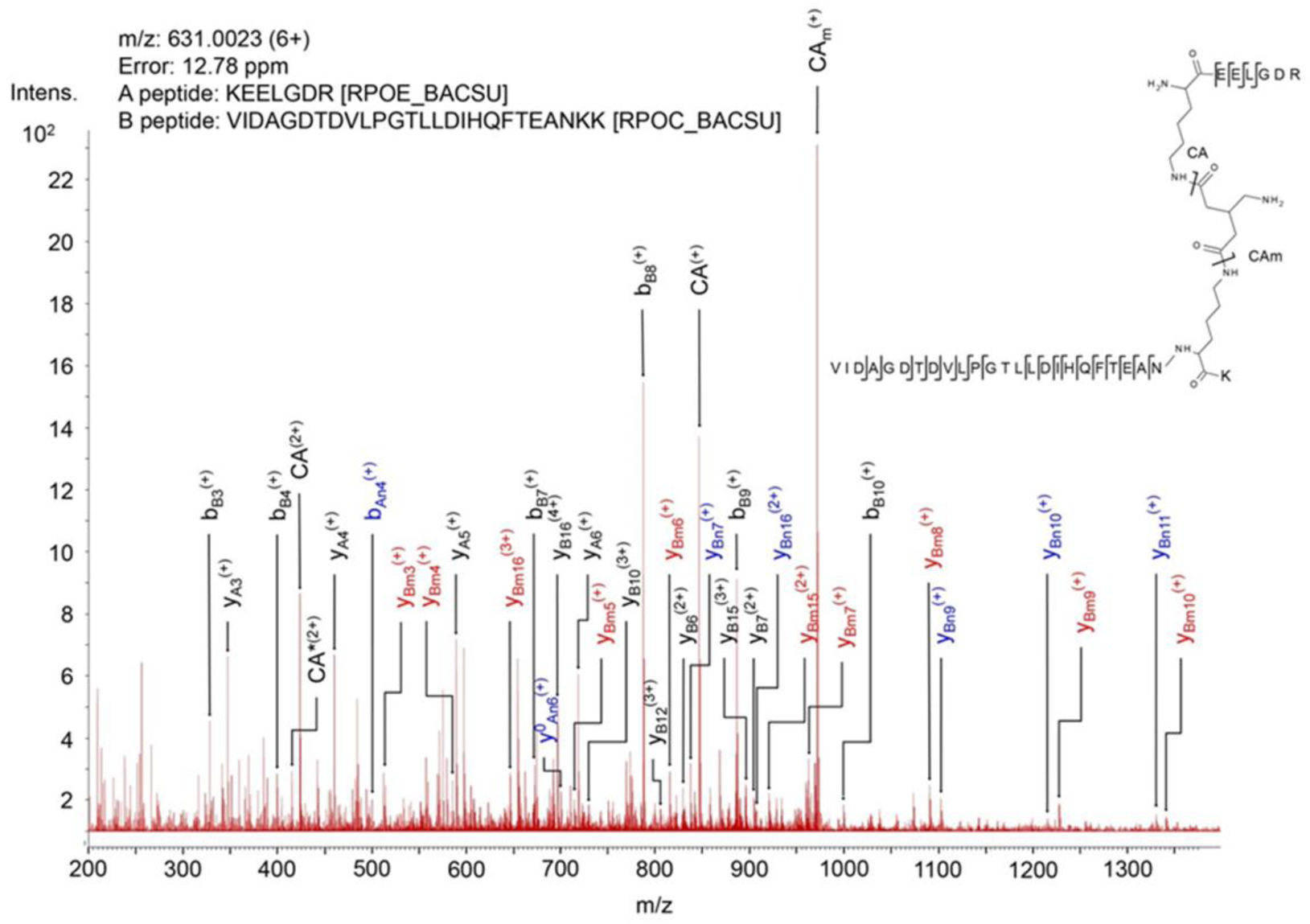
Mass spectrum of product ions generated by collision induced dissociation of a precursor ion of a BAMG-cross-linked peptide pair. The spectrum shows characteristic features of the fragmentation pattern of a cross-linked peptide in which the azido group in the spacer of the cross-linker has been reduced to an amine group. These features are (i) signals of high intensity resulting from cleavage of the cross-linked amide bonds leading to unmodified peptide A (CA) and peptide A modified by the remnant of the cross-linker in the form of a γ-lactam (CAm), adding 125.0477 Da to the mass of peptide and (ii) secondary fragments resulting from cleavage of a cross-linked amide bond along with peptide bond cleavages of an unmodified peptide (blue, subscript An, Bn) or a peptide lactam (red, subscript Am, Bm). These secondary cleavages occur along with primary cleavages of the peptide bonds (black, subscript A, B). The presence of both primary fragments (resulting from cleavages of the cross-link amide bonds and peptide bonds) and secondary fragments tremendously facilitates identification of cross-linked peptides according to the work flow schematically depicted in Fig. 3b. *, fragment with NH3 loss.

**Figure 5.**
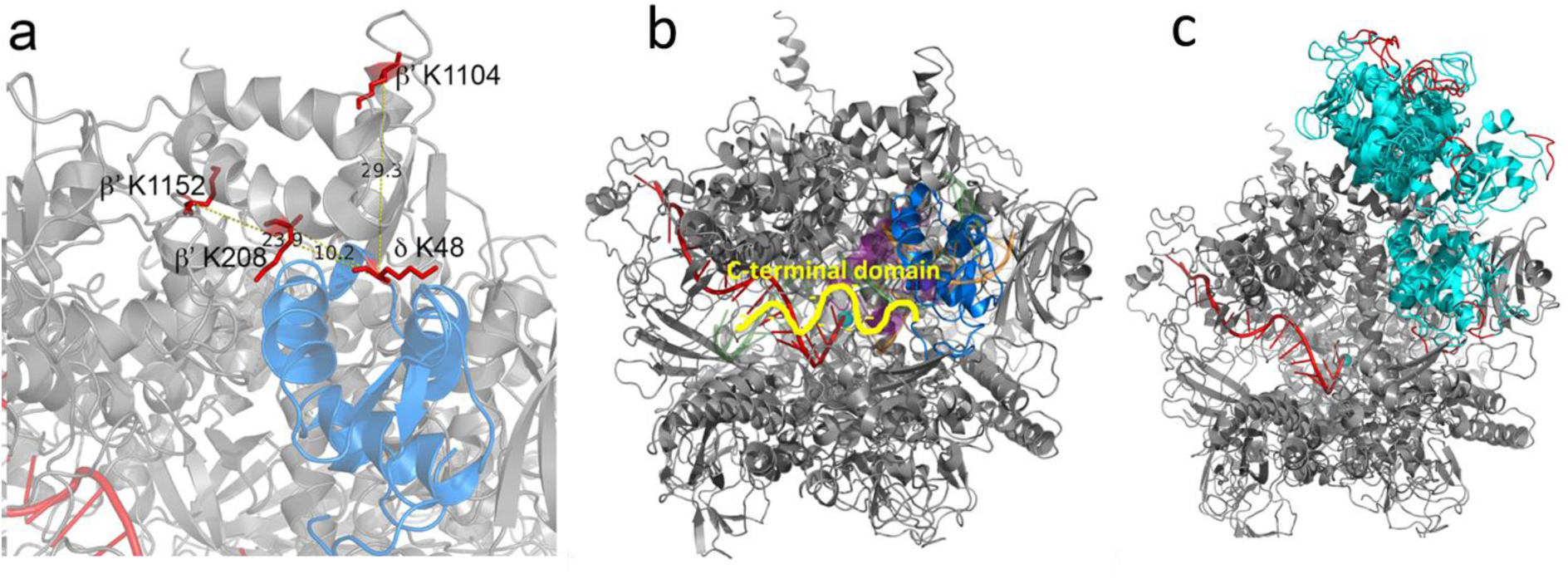
Model of *B. subtilis* RNAP in complex with δ. **a**, a zoomed region of δ (blue) located within the DNA binding cleft of RNAP (grey). The cross-linked amino acids are shown in red and the distances in Å between the cα carbon of δ K48 and RNAP β’ K208, K1104 and K1152 indicated. **b**, a model of RNAP (grey) in complex with δ (blue) with σ region 1.1 (purple) and DNA (green, template strand; orange, non-template strand) shown as semi-transparent cartoons. The active site Mg^2+^ is shown as a cyan sphere and RNA as a red cartoon. (Part of) the unstructered C-terminal domain, attached at the C-terminal end of the structured N-terminal domain, is depicted as a yellow squiggle pointing in the direction of the RNA export channel. **c**, compilation of all 10 docked models (all cyan) with the C-terminal 5 amino acids of the structured N-terminal domain colored red.

### Mass spectrometric analysis reveals a large number of cross-linked peptides at a low false discovery rate

The work-flow for sample preparation of cross-linked peptides from *in vivo* cross-linked *B. subtilis* cells for LC-MS/MS analysis is shown in Figure 2b. After cross-linking and protein extraction, cross-linked proteins were fractionated by size exclusion chromatography to obtain a sample expected to be enriched in cross-links formed during transient interaction. To this end samples with a size distribution of roughly 400 to 2000 kDa were used for further analysis. This fraction was enriched in RNAP and also contained ribosomes, i.e., protein complexes involved in processes guided by many transient protein-protein interactions. A list of proteins present in this fraction, identified by peptide fragment fingerprinting, and sorted according to their abundance index (29), is presented in Table 1 (Supporting Information). Besides subunits from ribosomes and RNAP, we also detected many proteins of high abundance with a known molecular weight far below 400 kDa, that included all glycolytic and TCA cycle enzymes, and many enzymes involved in amino acid synthesis, indicating that these proteins interact *in vivo* with other proteins.

**Table 1.**
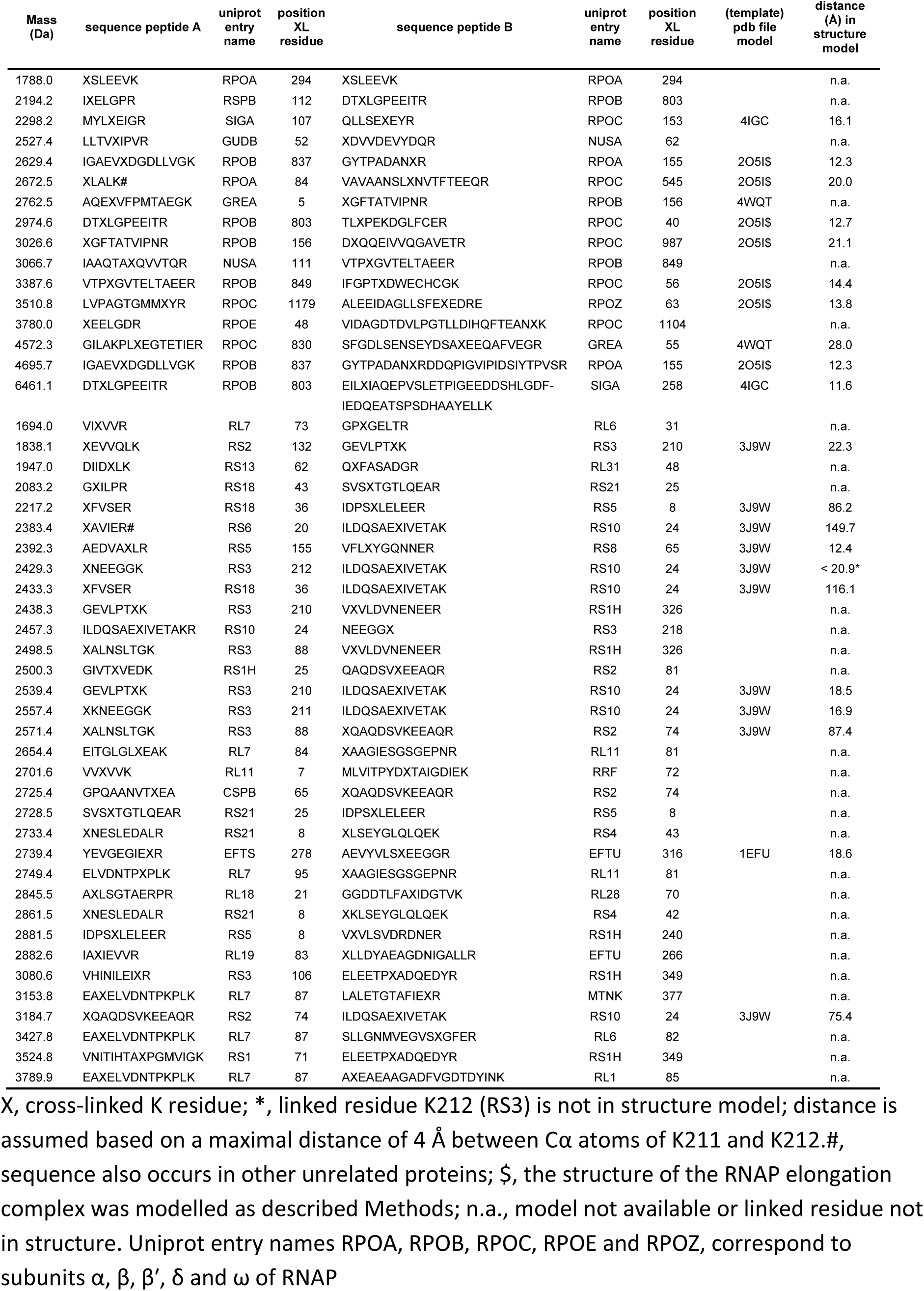
Inter-protein cross-linked peptides from proteins involved in transcription and translation

| Mass (Da) | sequence peptide A | uniprot entry name | position XL residue | sequence peptide B | uniprot entry name | position XL residue | (template) pdb file model | distance (Å) in structure model |
| --- | --- | --- | --- | --- | --- | --- | --- | --- |
| 1788.0 | XSLEEVK | RPOA | 294 | XSLEEVK | RPOA | 294 |  | n.a. |
| 2194.2 | IXELGPR | RSPB | 112 | DTXLGPPEITR | RPOB | 803 |  | n.a. |
| 2298.2 | MYLXEIGR | SIGA | 107 | QLLSEXEYR | RPOC | 153 | 4IGC | 16.1 |
| 2527.4 | LLTVXIPVR | GUDB | 52 | XDVVDEVYDQR | NUSA | 62 |  | n.a. |
| 2629.4 | IGAEVXDGDLLVGK | RPOB | 837 | GYTPADANXR | RPOA | 155 | 2O5I\$ | 12.3 |
| 2672.5 | XLALK# | RPOA | 84 | VAVAANSLXNVTFTEEQR | RPOC | 545 | 2O5I\$ | 20.0 |
| 2762.5 | AQEXVFPMTAEGK | GREA | 5 | XGFTATVIPNR | RPOB | 156 | 4WQT | n.a. |
| 2974.6 | DTXLGPPEITR | RPOB | 803 | TLXPEKDGFLFCER | RPOC | 40 | 2O5I\$ | 12.7 |
| 3026.6 | XGFTATVIPNR | RPOB | 156 | DXQQEIVVQGAVETR | RPOC | 987 | 2O5I\$ | 21.1 |
| 3066.7 | IAAQTAQVVTQR | NUSA | 111 | VTPXGVTeltaER | RPOB | 849 |  | n.a. |
| 3387.6 | VTPXGVTeltaER | RPOB | 849 | IFGPTXDWECHCGK | RPOC | 56 | 2O5I\$ | 14.4 |
| 3510.8 | LVPAGTGMMXYR | RPOC | 1179 | ALEEIDAGLLSFEXEDRE | RPOZ | 63 | 2O5I\$ | 13.8 |
| 3780.0 | XEELGDR | RPOE | 48 | VIDAGDTDVLPGLTDIHFTEANXK | RPOC | 1104 |  | n.a. |
| 4572.3 | GILAKPLXEGTETIER | RPOC | 830 | SFGDLSSENSEYDSAXEEQAFVEGR | GREA | 55 | 4WQT | 28.0 |
| 4695.7 | IGAEVXDGDLLVGK | RPOB | 837 | GYTPADANXRDDQPIGVIPIDSIYTPVSR | RPOA | 155 | 2O5I\$ | 12.3 |
| 6461.1 | DTXLGPPEITR | RPOB | 803 | EILXIAQEPVSLPTIGEEDDSHLGDF-<br>IEDQEATSPSDHAAYELLK | SIGA | 258 | 4IGC | 11.6 |
| 1694.0 | VIXVVR | RL7 | 73 | GPXGELTR | RL6 | 31 |  | n.a. |
| 1838.1 | XEVVQLK | RS2 | 132 | GEVLPTXK | RS3 | 210 | 3J9W | 22.3 |
| 1947.0 | DIIDLK | RS13 | 62 | QXFASADGR | RL31 | 48 |  | n.a. |
| 2083.2 | GXILPR | RS18 | 43 | SVSXTGTLQEAR | RS21 | 25 |  | n.a. |
| 2217.2 | XFVSR | RS18 | 36 | IDPSXLELEER | RS5 | 8 | 3J9W | 86.2 |
| 2383.4 | XAVIER# | RS6 | 20 | ILDQSAEXIVETAK | RS10 | 24 | 3J9W | 149.7 |
| 2392.3 | AEDVAXLR | RS5 | 155 | VFLXYGQNNER | RS8 | 65 | 3J9W | 12.4 |
| 2429.3 | XNEEGGK | RS3 | 212 | ILDQSAEXIVETAK | RS10 | 24 | 3J9W | < 20.9* |
| 2433.3 | XFVSR | RS18 | 36 | ILDQSAEXIVETAK | RS10 | 24 | 3J9W | 116.1 |
| 2438.3 | GEVLPTXK | RS3 | 210 | VXVLDVNEEER | RS1H | 326 |  | n.a. |
| 2457.3 | ILDQSAEXIVETAKR | RS10 | 24 | NEEGGX | RS3 | 218 |  | n.a. |
| 2498.5 | XALNSLTGK | RS3 | 88 | VXVLDVNEEER | RS1H | 326 |  | n.a. |
| 2500.3 | GIVTXVEDK | RS1H | 25 | QAQDSVKEEAQR | RS2 | 81 |  | n.a. |
| 2539.4 | GEVLPTXK | RS3 | 210 | ILDQSAEXIVETAK | RS10 | 24 | 3J9W | 18.5 |
| 2557.4 | XKNEEGGK | RS3 | 211 | ILDQSAEXIVETAK | RS10 | 24 | 3J9W | 16.9 |
| 2571.4 | XALNSLTGK | RS3 | 88 | XQAQDSVKEEAQR | RS2 | 74 | 3J9W | 87.4 |
| 2654.4 | EITGLGLXEA | RL7 | 84 | XAAGIESGSGEPNR | RL11 | 81 |  | n.a. |
| 2701.6 | VXVVK | RL11 | 7 | MLVITPYDXTAIGDIEK | RRF | 72 |  | n.a. |
| 2725.4 | GPQAANVTXEA | CSPB | 65 | XQAQDSVKEEAQR | RS2 | 74 |  | n.a. |
| 2728.5 | SVSXTGTLQEAR | RS21 | 25 | IDPSXLELEER | RS5 | 8 |  | n.a. |
| 2733.4 | XNESLEDALR | RS21 | 8 | XLSEYGLQLQEK | RS4 | 43 |  | n.a. |
| 2739.4 | YEVGEGIEXR | EFTS | 278 | AEVYVLSXEEGGR | EFTU | 316 | 1EFU | 18.6 |
| 2749.4 | ELVDNTPXPLK | RL7 | 95 | XAAGIESGSGEPNR | RL11 | 81 |  | n.a. |
| 2845.5 | AXLSGTAERPR | RL18 | 21 | GGDDTLFAXIDGTVK | RL28 | 70 |  | n.a. |
| 2861.5 | XNESLEDALR | RS21 | 8 | XKLSEYGLQLQEK | RS4 | 42 |  | n.a. |
| 2881.5 | IDPSXLELEER | RS5 | 8 | VXVLSVDRDNER | RS1H | 240 |  | n.a. |
| 2882.6 | IAXIEVVR | RL19 | 83 | XLLDYAEAGDNIGALLR | EFTU | 266 |  | n.a. |
| 3080.6 | VHINILEIXR | RS3 | 106 | ELEETPXADQEDYR | RS1H | 349 |  | n.a. |
| 3153.8 | EAXELVDNTPKPLK | RL7 | 87 | LALETGTAFIEXR | MTNK | 377 |  | n.a. |
| 3184.7 | XQAQDSVKEEAQR | RS2 | 74 | ILDQSAEXIVETAK | RS10 | 24 | 3J9W | 75.4 |
| 3427.8 | EAXELVDNTPKPLK | RL7 | 87 | SLLGNMVEGVXGFER | RL6 | 82 |  | n.a. |
| 3524.8 | VNITIHTAXPGMVGK | RS1 | 71 | ELEETPXADQEDYR | RS1H | 349 |  | n.a. |
| 3789.9 | EAXELVDNTPKPLK | RL7 | 87 | AXEAEAGADFGDIDYINK | RL1 | 85 |  | n.a. |

3 X, cross-linked K residue; \*, linked residue K212 (RS3) is not in structure model; distance is 4 assumed based on a maximal distance of 4 Å between Cα atoms of K211 and K212. #, 5 sequence also occurs in other unrelated proteins; \$, the structure of the RNAP elongation 6 complex was modelled as described Methods; n.a., model not available or linked residue not 7 in structure. Uniprot entry names RPOA, RPOB, RPOC, RPOE and RPOZ, correspond to 8 subunits α, β, β', δ and ω of RNAP

After trypsin digestion of the extracted proteins in the high molecular weight fraction, cross-linked peptides were enriched by diagonal strong cation exchange (SCX) chromatography (9).The principle of the enrichment is schematically depicted in Figure 2b. Peptides in the cross-link-enriched SCX fractions were subjected to LC-MS/MS, data processing and database searching, according to the workflow schematically depicted in Figure 3 (10). For efficient identification of cross-linked peptides from the entire *B. subtilis* sequence database with MS/MS data, it is necessary to know the masses of the two peptides in a cross-link. This is possible due to abundant signals in MS/MS spectra arising from cleavage of the two cross-linked amide bonds (10) shown as an example in the mass spectrum in Figure 4. Following this protocol, we identified 82 unique inter-protein cross-links (Table 2, Supporting Information) and 369 unique intra-protein cross-links (Table 3, Supporting Information) in 299 and 1920 precursor ions, respectively, that fulfilled all criteria mentioned in Figure 3. Importantly, no decoy peptides fulfilled these criteria, indicating a very low false discovery rate (FDR).

About 39% of the 82 unique inter-protein cross-linked peptides are from enzymes involved in intermediary metabolism, protein and RNA folding, and protein and RNA degradation. Most of these cross-links comprise peptides with identical sequences, showing that the parent proteins occurred in symmetric homodimers, possibly organized in higher order assemblies. About 40% of all inter-protein cross-links are from translation complexes, i.e., ribosomes and auxiliary proteins involved in translation, and about 18% are from transcription complexes, i.e., RNAP and initiation and elongation factors (Table 2, Supporting Information).

### Use of different assignment criteria for inter-protein and intra-protein cross-linked peptides

To obtain a low FDR of both intra-protein and inter-protein cross-links we used different assignment criteria for these two types of cross-links (Figure 3). Since the probability of identifying false positive cross-linked peptide pairs from a complete sequence database is much higher for inter-protein cross-links than for intra-protein cross-links, false discoveries are practically all confined to inter-protein cross-links if the same assignment criteria are employed for both cross-link type (10,21,22). In Table 4 (Supporting Information) it is shown how variations in assignment criteria affect the number of identified cross-links and the FDR. Applying the more stringent criteria for assignment of inter-protein cross-links to intra-protein cross-links only leads to a decrease of approximately 20% assigned unique cross-linked peptides. Consequently, the number of assigned inter-protein cross-links slightly increases upon relaxing the stringency of the criteria for assignment. However, this increase is accompanied by a relatively large increase in FDR. So, the stringent criteria that we apply here for inter-protein cross-links result in efficient identification and an extremely low FDR.

### Biological consistency of identified cross-linked peptides

To corroborate identified cross-linked peptides by comparison with published data we determined spatial distances between Cα atoms of linked residues. In models of crystal structures, the maximal distance that can be spanned by BAMG varies between 25.7 to 29.7 Å, assuming a spacer length of BAMG of 7.7 Å, a lysine side chain length of 6.5 Å and a coordinate error of 2.5 – 4.5 Å. The distances between 95.6% (n = 135) of Cα atom pairs of linked residues in cross-links with non-overlapping sequences from one protein (denoted intra-protein cross-links) are less than 25.7 Å, including 14 inter-protein cross-links between identical proteins that fitted better than intra-protein species (Table 3 and Figure 6 of the Supporting Information). The distances between Cα atoms of linked residues of only 2 cross-links out of the 135 exceed 29.7 Å. These results underscore the high biological consistency and, thereby, reliability of identified cross-linked peptides.

Table 1 lists the inter-protein cross-linked peptides from transcription and translation complexes. The distances between the Cα atoms of interlinked lysine residues of all 9 cross-links comprising peptides from proteins involved in transcription are in agreement with models based on crystal structures. Also 5 small ribosomal inter-protein cross-linked peptides nicely fit in the available structural model of a stalled ribosome (30). However, five small ribosomal inter-protein cross-links that exceed 29.7 Å by more than 45 Å were notable exceptions. Since the FDR is extremely low and the large majority of our dataset is biologically consistent, it is reasonable to assume that formation of these cross-links actually took place. Most likely these distance measurements represent the detection of ribosomal assembly intermediates and/or covalent links between juxtaposed ribosomes.

### Many cross-links reveal transient protein-protein interactions

The power of our approach was demonstrated by the detection of several transient interactions between translation factors and ribosomes and between transcription factors and core RNAP (Table 1). Ribosome-recycling factor RRF forms a cross-link with ribosomal protein RL11, in agreement with cryo-EM data showing an interaction between these two proteins in the post-termination complex (31). A cross-link between RL19 and EF-Tu is in agreement with the presence of RL19 near the EF-Tu binding site on the ribosome (32). Cross-linked peptides were found between K4 and K55 of the transcription elongation factor GreA and residues β-K156 and β'-K830, respectively, in the RNAP secondary channel. This position fits with the known function of GreA and with a crystal structure of a chimeric Gfh1-GreA in complex with RNAP (33). Likewise, the binding of NusA close to the RNA exit channel of RNAP, as revealed by a cross-link between NusA-K111 and β-K849, is in agreement with results obtained previously that indicates the N-terminal domain of NusA binds to the β-flap tip of RNAP (24,34). Two cross-linked peptides between the sigma A factor (σ^A^) and RNAP were identified. The distances between Cα atoms of corresponding residues in the structure of the *E. coli* RNAP holoenzyme (35) are 16.1 Å and 11.6 Å. Thus, the spatial arrangements of the proteins involved in these transient interactions, are in agreement with previously published *in vitro* data, underscoring the reliability of our *in vivo* cross-link approach.

Of great interest was the identification of novel transient interactions. A binding site of the RNA chaperone CspB on ribosomes, as revealed by a cross-link between CspB and RS2, has not been observed before to our knowledge. This interaction makes sense, since cold shock proteins co-localise with ribosomes in live cells and are involved in coupling transcription and translation (36,37). The biological significance of the interaction between glutamate dehydrogenase GudB and transcription elongation factor NusA is not known, but recent work may suggest a functional link between the two proteins. The *gudB* gene encodes a cryptic glutamate dehydrogenase (GDH), which is highly expressed but not active. If the main GDH (RocG) is inactivated, a frame-shift mutation activates GudB. This mutation depends on transcription of *gudB*, and requires the transcription-repair coupling factor Mfd (38). Interestingly, NusA is also involved in transcription-coupled repair (39). Whether the interaction of GudB with NusA is relevant for the regulation of this gene decryption remains to be established.

Another noteworthy interaction is revealed by a cross-link between the βʹ subunit of RNAP and a protein originally found associated with isolated RNAP named δ (40). Importantly, δ has a complex effect on transcription. It inhibits initiation from weak promoters mediated by σ^A^ (41,42), stimulates or inhibits transcription from certain other promotors (43,44) and increases RNAP recycling speed (42,43) in synergy with the DNA helicase HelD (45), probably by dissociation of stalled RNAP-DNA or RNAP-RNA complexes. Transcriptomics experiments indicate δ reduces non-specific initiation of transcription which is relatively prevalent in Gram positive bacteria (46,47). Up to now it has remained elusive how these different effects on transcription are brought about.

The 20.4 kDa δ occurs exclusively in Gram-positive bacteria. It consists of an amino-terminal globular domain and a nucleic acid-mimicking highly acidic unstructured C-terminal half (48,49). The δ protein forms a complex with the RNAP core enzyme in a 1:1 stoichiometry (43,48,50). A truncated form of δ lacking the C-terminal half is sufficient for binding to RNAP. Intact δ, as well as the acidic unstructured C-terminal domain, but not the truncated N-terminal domain inhibits the binding of nucleic acids to RNAP. It has been proposed that the amino-terminal RNAP-binding domain may act both to orient and increase the local concentration of the flexible negatively charged carboxyl terminal domain to effectively shield nearby positively charged nucleic acid binding sites on RNAP (48). The effect of δ on promoter selectivity by σ^A^-mediated initiation suggests an interaction of δ with the preinitiation complex. Indeed, it has been reported that δ and σ^A^ can bindsimultaneously to core RNAP, with negative cooperativity (51). However, other experiments indicated mutual exclusion of the binding of δ and σ^A^ to core RNAP (43,52).

We identified a cross-link between K48 of the δ subunit (RpoE) and K1104 of the RNAP β' subunit (RpoC) (Figure 4). K1104 is located in the so-called downstream clamp region. This suggests a binding site for the δ subunit on RNAP close to the downstream DNA binding cleft. To confirm this finding, we performed *in vitro* cross-linking with purified δ-containing RNAP. This resulted in two additional cross-links, one between K48 of δ and residue β'-K208, and one between δ-K48 and β'-K1152, both in close proximity to β'-K1104, thereby corroborating our *in vivo* findings. Since only one residue in the N-terminal domain of δ was involved in cross-linking, further evidence is required to assign a preferential orientation of delta with respect to the clamp region. To this end we used in an *in silico*docking analysis using a *B. subtilis* RNAP elongation complex model and the known N-terminal domain structure of δ (24,53). The best 10 output models were analyzed to establish which complied with the maximum Cα-Cα cross-link distance achievable with BAMG (Table 5, Supporting Information). In all but one model, at least one cross-link exceeded the 29.7 Å maximum distance, however, in published structures of RNAP, crystallographic B factors are relatively high around positions β’-K1104 and β’-K1152, implying some conformational flexibility in those regions. The model which gave the lowest cumulative Cα-Cα cross-link distance, with all predicted cross-links < 29.7 Å places δ in the downstream side of the DNA binding cleft of RNAP (Figure 5a). A position of δ inside the DNA binding cleft as shown in Figure 5b suggests that the N-terminal domain of δ could sterically inhibit the binding of downstream DNA. In this position δ could also sterically inhibit the binding of the 1.1 region of σ^A^, since this region is expected to interact with an overlapping binding site, based on crystal structures of *E. coli* RNAP holoenzyme with the homologous σ^70^ factor (27,35). The model with the next lowest aggregate score also placed δ in this region, but the remaining 8 placed it on top of the β’ subunit outside of the DNA binding cleft (Figure 5c). This position of the N-terminal domain of δ outside the downstream DNA channel but close to its entrance implies that interference with DNA binding and binding of the 1.1 region of the σ factor requires penetration of the C-terminal unstructured acid domain into the channel to interact with positive charges of the polymerase involved in DNA binding and in binding of the σ 1.1 region. Since δ is known to displace RNA more efficiently than it can DNA (48) and is important in RNAP recycling following the termination of transcription (45), we expect that it must be oriented so that the acidic C-terminal domain is able to influence RNA binding through contact with the transcript close to or even within the RNA exit channel. However, the data prevent discrimination between models placing δ inside or outside the downstream DNA channel, since both the aggregate distance scores (Table 5, Supporting Information) and the HADDOCK scores of the best models show only small differences.

## DISCUSSION

We have developed a new method for proteome-wide identification of cross-links introduced *in vivo* by N-hydroxysuccinimidyl esters directly in a bacterial cell culture. Within as little as 5 min extensive cross-linking was observed, using 2 mM BAMG. This implies that cross-link analysis on a time scale of seconds could be a future development, enabling, for instance, monitoring at the peptide level transient protein-protein interactions involved in rapid cellular adaptation. Rapid in vivo cross-linking requires a Gram-positive organism, since the outer membrane of Gram-negative species forms a barrier for diffusion.

With our method we identified many inter-protein cross-links confirming several known stable and transient protein interactions, underscoring the reliability of the approach. In addition, we found intriguing new interactions that deserve further investigation to understand their functional significance, like the interaction between GudB and NusA. Another intriguing cross-link was found between the δ and βʹ subunits of RNAP. This cross-link revealed that δ binds to the clamp region of RNAP close to the entrance of the downstream DNA binding cleft. However, the cross-link data combined with *in silico* docking experiments did not provide enough evidence to assign an unambiguous orientation of δ with respect to the channel. This implies that different scenarios are possible for the molecular mechanism by which δ regulates promoter selectivity and promotes RNAP recycling. However, the knowledge that δ binds to the clamp region of βʹ will help to design experiments aimed at developing a better understanding of the mechanism of action of δ.

Besides membrane permeability, the unique chemical properties of BAMG, combined with our statistical analysis, are at the heart of the large number of identified cross-linked peptides with high biological consistency and low false discovery rate. This is also the first study in which a large number of cross-links from intracellular protein complexes generated in undisturbed growing cells have been identified by mass spectrometry and database searching using a complete species specific database. Our dataset consists of about 18% inter-protein and 82% intra-protein cross-links. This percentage of inter-protein cross-links is relatively low in comparison with other datasets obtained either by *in vivo* cross-linking of bacteria (2-5) or by cross-linking bacterial cellular protein extracts of high complexity (54,55). Also, the reported fractions of inter-protein cross-links identified in human cells, 23-25% of the total number of cross-links (56,57), is relatively high compared with the 10% that we obtained in a previous study (10). We consider it important for the future development of the technology to understand the causes of such differences. In one of these studies (57) we noticed that many sequences of the peptides from identified cross-links, classified as intermolecular species, are not unique, and therefore that many cross-linked peptide pairs could either be from the same protein or from different proteins. Furthermore, in protein extracts non-specific interactions may be formed dependent on extraction conditions. However, a key difference between these reports (2-5,54-57) and our approach concerns the statistical analysis, in which we make a distinction between inter-and intra-molecular cross-links. If this distinction is not made, an overestimation of intermolecular cross-links occurs at the expense of a relatively large number of false positives.

In this study we provide an efficient layout for proteome scale crosslink identification using in-culture crosslinking. Several modifications to the method can be made to tailor specific experimental requirements. For example, in proteome-wide analyses low abundance cross-links can escape detection by LC-MS/MS. The use of affinity tagged target proteins will enable enrichment of these complexes for subsequent inter-peptide cross-link identification of transient interactions. Furthermore, in this study we have focused on the soluble fraction of cross-linked cells. Further extraction and digestion of the insoluble fraction, enriched in membrane and cell wall proteins, is likely to reveal a rich source of interesting protein crosslinks. Relative quantification of cross-linked peptides can also be employed with commercially available isotope-labeled starting materials for the synthesis route of BAMG presented here. Finally, our analytical strategy may also benefit from the option of mass spectrometry to combine collision-induced dissociation with electron transfer dissociation (57,58) to increase efficiency of identification of cross-linked peptides. The high average precursor charge state of slightly more than +4 of all identified BAMG-cross-linked peptides in our dataset is favorable for the latter fragmentation method (59). Overall, we believe that the *in vivo* cross-linking and data analysis methods developed here will pave the way to a systems level view on dynamic protein interactions. Such a view will lead to a deeper understanding of the molecular mechanisms of biological processes guided by dynamic protein-protein interactions in the cell.

## Conclusions

A system has been developed for rapid *in vivo* protein cross-linking by an amine-specific bifunctional reagent added directly to a culture of *Bacillus subtilis.* We identified several stable and dynamic interactions in intracellular protein complexes with a size range of about 400 to 2000 kDa by mass spectrometric analysis of isolated cross-linked peptides and database searching using the entire species specific sequence database. In culture cross-linking of cytoplasmic protein in Gram-positive bacteria is much faster than in Gram-negative species. In combination with affinity purification of target proteins, our *in vivo* cross-link technology will be useful to obtain insight in the molecular mechanisms of processes guided by dynamic protein-protein interactions in large assemblies.

## Acknowledgements

Work in P.J.L’s laboratory was supported by a grant from the Australian Research Council (ARC, DP110100190)

## Supporting Information

PDF files: Figures S1-S6; Table S4-S5

Excel files: Table S1-S3

## Additional Information

The programs REANG and YEUN YAN, written in Visual Basics for Applications will be send on request by the authors W.R: or H.B

